# Multiple independent components contribute to event-related potential correlates of conscious vision

**DOI:** 10.1101/2023.01.02.522455

**Authors:** Elisabetta Colombari, Henry Railo

## Abstract

Research has revealed two major event-related potential (ERP) markers of visual awareness: the earlier Visual Awareness Negativity (VAN, around 150–250 ms after stimulus onset), and the following Late Positivity (LP, around 300–500 ms after stimulus onset). Understanding the neural sources that give rise to VAN and LP is important in order to understand what kind of neural processes enable conscious visual perception. Although the ERPs afford high temporal resolution, their spatial resolution is limited because multiple separate sources sum up at the scalp level. In the present study, we sought to characterize the locations and time-courses of independent neural sources underlying the ERP correlates of visual awareness by means of Independent Component Analysis (ICA). ICA allows identifying and localizing the temporal dynamics of different neural sources that contribute to the ERP correlates of conscious perception. The present results show that while LP reflects a combination of multiple sources distributed among frontal, parietal and occipito-temporal cortex, the sources of VAN are localized to posterior areas including occipital and temporal cortex. In addition, our analysis reveals that activity in very early sources (roughly -100–100 ms after stimulus onset) in temporal and fronto-parietal cortices correlates with conscious vision.

## 1. Introduction

Over the past years, great efforts have been devoted to the search for the neural correlates of consciousness (NCC). One of the main lines of study searches for the NCC that underlies the emergence of conscious visual experience. Due to their high temporal resolution, event-related potentials (ERP) measured using electroencephalography (EEG) provide excellent means to examine this question (Luck, 2014). These studies have revealed scalp recorded electrophysiological signatures of conscious vision, but the neural processes that generate these correlates are not well understood. Here, we sought to shed light on the spatio-temporal distribution of neural sources that contribute to the ERP correlates of conscious vision.

The most widely used approach to investigate the NCC consists of presenting participants with a stimulus that they only sometimes consciously perceive. This allows contrasting the brain activity associated with subjectively “seen” and “unseen” stimuli while keeping the objective, physical stimulus constant. This procedure allows detecting the neural processes that are involved in visual awareness, although processes that are not specifically linked with conscious vision are likely also present (Aru *et al*., 2012). Processes that co-vary with conscious perception, but are not likely part of the mechanism that directly enables conscious perception can be roughly categorized into two classes: processes that contribute to perception, but take place before conscious perception (e.g., prestimulus activity (Britz *et al*., 2014)), and later processes that are caused by visual conscious perception (e.g., cognitive processes and behavioral response-related processes).

The comparison between the Aware (i.e., “seen”) and Unaware (i.e., “unseen”) conditions has revealed two major ERP correlates of conscious vision: a negative amplitude difference, typically peaking around 200 ms after the stimulus onset in occipito-temporal sites, called Visual Awareness Negativity (VAN), and a later enhanced positivity (i.e., Late Positivity, LP) occurring at centro-parietal electrodes in the P3 time window (i.e., around 300-500 ms after the presentation of the stimulus) (Koivisto & Revonsuo, 2003, 2010; Dehaene & Changeux, 2011; Mashour *et al*., 2020). While VAN is known to be localized over occipito-temporal sites (Veser *et al*., 2008; Koivisto & Revonsuo, 2010; Liu *et al*., 2012), the topography of LP is widely distributed over multiple cerebral sources spanning occipital, parietal, temporal, and frontal cortices. Furthermore, LP is thought to reflect several different cognitive processes such as identifying, naming and reporting the stimulus (Volpe *et al*., 2007; Koivisto & Revonsuo, 2010; Dehaene & Changeux, 2011). Accordingly, the localization of the NCC is still unclear. In particular, debate continues over how crucial frontal areas are for conscious perception: because of its involvement in cognitive functions such as attention and working memory, the prefrontal cortex is sometimes argued to play an essential role in conscious perception (Del Cul et al., 2009, Odegaard et al., 2017). In contrast, other evidence suggests that frontal and prefrontal cortices may be neither necessary nor sufficient for consciousness (Boly *et al*., 2017; Raccah *et al*., 2021), suggesting that the NCC are localized in posterior cortical regions, including occipital, parietal and temporal lobes (Koch et al., 2016; Mazzi & Savazzi, 2019). Given that ERPs are the summed activity of multiple distinct sources, correlates such as VAN and LP may at a given time-point include a combination of consciousness-related sources, instead of being localized to a single area. Moreover, because sources with opposite polarities may cancel each other out, they may become invisible in the average ERP (Luck & Kappenman, 2012). Theories about the neural basis of consciousness typically argue that conscious perception involves recurrent activity across multiple areas, suggesting that correlates such as VAN and LP are a combination of multiple sources. This is especially true for LP, which according to the Global Neuronal Workspace Theory of consciousness, involves co-organized activity across widely distributed cortical areas (Dehaene & Changeux, 2011). While early correlates are typically argued to be driven by posterior areas, it remains possible that early activity in frontal areas also correlates with consciousness if a sufficiently sensitive source separation is utilized (Thompson & Schall, 2000; Knotts *et al*., 2018; Kapoor *et al*., 2022).

During recent years, source separation approaches such as independent component analysis (ICA) have been developed to uncover sources contributing to average ERPs. In ICA, EEG is decomposed into maximally independent components (ICs) (Onton & Makeig, 2006). Each IC represents a temporally and functionally independent source of the EEG signal, with a specific scalp distribution (which is constant over time), and a specific amplitude at each time point (Onton & Makeig, 2006; Onton et al., 2006). This allows investigation of source level activity and isolating ICs that underlie the average ERP wave.

The aim of the present study was to characterize the locations and time-courses of independent neural sources that significantly contribute to the ERP correlates of visual awareness. We expected to identify components that significantly contribute to VAN and LP, but hypothesized that source separation might also uncover correlates that are not visible in scalp recorded ERPs.

## 2. Materials and Methods

### 2.1 Participants

Analyses presented in this study were performed on data acquired in a previous study (Railo *et al*., 2021), and as part of an EEG course organized at the University of Turku (same paradigm and EEG methodology as in Railo et al., 2021). In the current study, data from 36 healthy participants (mean age ± sd = 24.14 ± 3.52, range 19-36) were analyzed. All participants were students at the University of Turku, and reported no neurological disorders. All of them gave their written informed consent in accordance with the Declaration of Helsinki. The study was approved by the ethics committee of the Hospital District of Southwest Finland.

### 2.2 Experimental procedure and Stimuli

The target stimuli were low contrast Gabor patches (diameter 6.5°, frequency 0.7 cycle/degree) which were presented to the left or right hemifield (about 5° from fixation on horizontal meridian) for 16.6 ms. In addition, the experiment included catch trials where no stimulus was presented (1/6^th^ of all trials). The intensity of the low contrast stimuli was individually determined using a QUEST staircase (Watson, 2017), so that stimulus intensity was near 50% of subjective detection threshold.

The stimulus was presented after a fixation period ranging from 668 to 1332 ms across trials. Participants were asked to report the location of the target (left vs. right), and to subjectively rate the visibility of the stimulus by means of a four-steps scale (Figure 1a). The scale was composed of the following four alternatives: 0) “did not see stimulus at all”, 1) “not sure but possibly saw something”, 2) “pretty sure I saw it”, 3) “saw the stimulus clearly”. A total of 400 trials was collected per participant (divided into 10 blocks of 40 trials).

**Figure 1.**
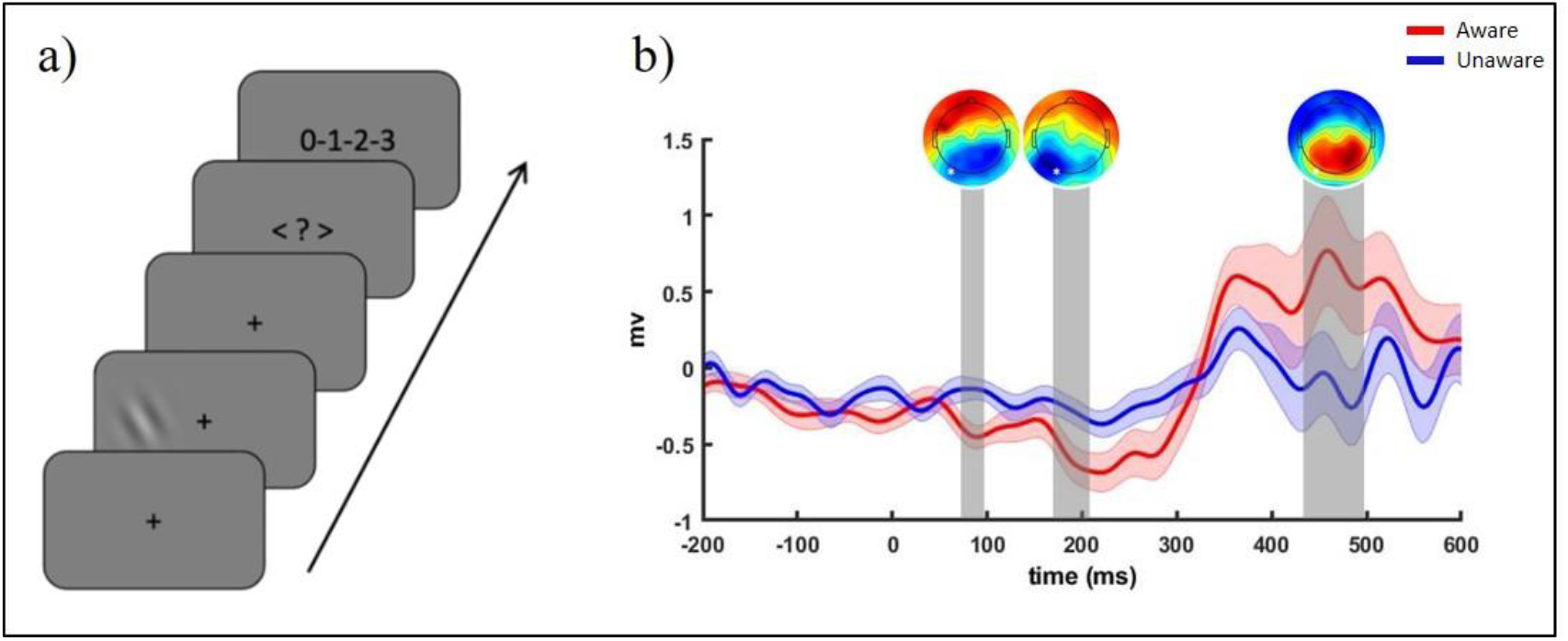
Experimental procedure and ERP results. A) Schematic presentation of a single experimental trial. B) Grand-average of ERPs computed for Aware and Unaware conditions at channel O1 (marked with a white star). Significant time-windows are highlighted in grey. The shaded area of the waveforms represents SEM at each time point, and scalp distribution maps represent the voltage difference between conditions

### 2.3 EEG

#### 2.3.1 Recording

EEG data were recorded with a 64-channel EEG system at a sampling rate of 500Hz. Impedance was kept near 5 kΩ. Electrode Fz served as on-line reference, and the ground electrode was placed on the forehead of the participant.

#### 2.3.2 Preprocessing

EEG data were preprocessed with MATLAB (version R2017b; the MathWorks, Inc., Natick, MA) using functions from the EEGLAB toolbox (v2020.0, Delorme & Makeig, 2004). Continuous raw data were first resampled to 250 Hz, and filtered using a high-pass filter at 1 Hz (cut-off frequency .5 Hz, transition bandwidth: 1Hz), followed by a low pass filter at 40 Hz (cut-off frequency 45Hz, transition bandwidth: 10Hz). *Clean_channels* EEGLAB function with a correlation threshold of .5 was used to remove channels with a bad signal (mean number of channels removed across participants = 1.61; SD = 1.84). After that, data were re-referenced to the average of all electrodes and cut into epochs ranging from -500 to 900 ms with respect to the stimulus onset. To remove epochs containing artefacts, ICA was computed using the extended Infomax runICA algorithm (Bell & Sejnowski, 1995), and trials contaminated by artefactual components were removed using the EEGLAB function *pop_jointprob* (SD = 5 for both local threshold and global threshold). Baseline correction was then applied on the pre-stimulus period (from -500 ms to 0 ms), and ICA was computed again.

Subsequently, the dipolar source of each component was localized using the DIPFIT plug-in (v3.3). The dipole localization was based on an average MRI, and electrode locations were co-registered based on standard channel coordinates. Because individual MRIs and subject-specific channel location information were not available, the accuracy of spatial localization in the present study is limited. Components with a residual variance of more than 15%, and those labelled as not-brain-based with a probability of >50% were automatically identified by means of the *ICLabel* plugin and removed. In total, 379 ICs (average number of ICs across participants =10.8) were selected as brain ICs. Finally, missing channels were interpolated for each participant using a spherical method using the EEGlab function *pop_interp*.

#### 2.3.3 IC clustering

The resulting 379 brain ICs were grouped into clusters using a k-means clustering method implemented in EEGlab. Components were clustered based on dipole location, dipole orientation and ERPs. Importantly, to avoid statistical “double-dipping” (Kriegeskorte *et al*., 2010), the data were divided into the Aware (visibility rating = 1,2,3) and Unaware (visibility rating = 0) experimental conditions only after the clustering. On average, the number of epochs included in the analyses was 107 (SD=35.92) for Aware and 105 (SD=38.18) for Unaware condition. The default number of clusters suggested by EEGlab (i.e., *k* value) was 11, but it was manually adjusted to 13 after visual inspection of the initial clustering result. On average, each cluster was composed of 35.2 ICs. Outliers ICs (35 ICs, threshold = 3 SD)—that is, ICs that were not assigned into any one of the 13 clusters— were grouped in an auxiliary cluster, which was not included in the analysis. For statistical analyses, ICs of each participant within a cluster were averaged together.

#### 2.3.4 Statistical analyses

Before IC clustering, we examined average ERP correlates of conscious vision, by comparing Aware (visibility ratings > 0) and Unaware conditions (rating = 0) using paired-samples t-tests computed on each time point. The p-values were corrected for multiple comparisons using Benjamini-Hochberg false-discovery rate procedure (Groppe *et al*., 2011) implemented in Matlab. Subsequently, for the analysis of IC correlates of conscious vision, the data were grouped into clusters, and within each cluster, ERPs were averaged separately for the two experimental conditions. Visual inspection of clusters suggested Aware vs Unaware differences resembling the VAN-LP pattern, including a transient early and later more sustained difference in many clusters. To maximize statistical power (and minimize the number of statistical tests), statistical analysis (FDR corrected paired-samples t tests) was focused on these time-windows (described in detail in the Results section). Since we were interested in quantifying the contribution of each cluster to the average ERP, we calculated for each cluster the percent variance accounted for (*pvaf*, which compares the variance of the whole data minus the back-projected component to the variance of the whole data) using the *std_envtopo* (v4.10) EEGlab function.

## 3. Results

The behavioral results showed that participants reported perceiving the stimulus on average in 48.27% of trials. Stimulus location discrimination accuracy was significantly greater for Aware trials (M = 95.02%) than Unaware trials (M = 64.91%; t(35) = 14.79, p < 0.001), indicating that in the Aware condition participants could properly discriminate the side of presentation of the stimulus. Mean RTs for Aware trials (1107.45 ms) were significantly faster than mean RTs for Unaware condition (1398.53 ms) (t(35) = -7.54, p < 0.001). In both conditions, RTs were longer than the ERP time window of interest (from -200 ms to 600 ms, with respect to the stimulus onset), meaning that the button press response should not contaminate the observed correlates of conscious vision.

Concerning the EEG results, as expected, both the VAN and the LP were observed in the grand average ERP. Figure 1b shows the grand average ERP for the Aware and Unaware conditions at electrode O1. Paired-sample t-test (FDR-corrected) between Aware and Unaware trials revealed that at electrode O1 VAN was significant in 2 different temporal windows: between 80 and 100 ms, and between 180 and 208 ms. LP was significant between 444 and 500 ms, although visual inspection suggests that its onset was earlier (around 325 ms).

To examine the independent brain sources that contribute to the scalp recorded ERPs, and possibly correlate with conscious vision, we next analyzed the clusters of ICs. Only clusters with more than 15 components were included into statistical analyses (clusters numbers 1, 2, 6, 7, 9, 10, 11, 12, 13). Excluded clusters contained on average 6 ICS (SD= 5.83). Figure 2 displays the ERP of each cluster, as well as the locations of individual ICs within the cluster. The clusters localized to different areas of the brain. For example, Cluster 1 localized to anterior parietal/posterior frontal areas, and Cluster 2 to temporal cortex. Visual inspection of the event-related responses suggested that especially in later time-windows (i.e., > 300 ms), the Aware and Unaware conditions differed. However, also earlier time-windows showed differences between the conditions, albeit smaller in amplitude (e.g., Clusters 2 and 12). Aware vs Unaware epochs were contrasted within each cluster in the selected time-windows (highlighted in Fig.2 by vertical lines) using FDR-corrected paired-sample t-tests. The results are reported in Table 1. The last column of the table shows the percent variance accounted for (pvaf) by each component at each temporal window. As shown in Table 1, many clusters showed statistically significant differences in the late time-windows (i.e., >300 ms after the stimulus presentation), and a few clusters also showed earlier differences. The earliest difference between Aware and Unaware conditions was observed -108–100 ms in parietal regions (Cluster 9). Also, Cluster 2 located in the temporal cortex showed a broad, early effect (32–212 ms), and also two later effects. Similar pattern is observed in Cluster 13 in the frontal cortex, although the difference between Aware and Unaware conditions was not statistically significant. Clusters 7 and 12 showed a mid-latency (roughly 150–260 ms) effect in occipito-temporal regions, in a time window corresponding to VAN in the scalp recorded grand average ERP. Finally, a later difference between Aware and Unaware conditions was observed in most of the clusters, spanning frontal, parietal, temporal and occipital regions.

**Table 1:** Results of FDR-corrected paired-sample t-tests computed contrasting Aware and Unaware conditions within each cluster in the reported time-windows.

| Cluster number | Temporal windows | p-value (fdr corrected) | Pvaf (%) |
| --- | --- | --- | --- |
| 1 | 348-556 | .0030* | 26.02 |
| 2 | 32-212 | .0101* | -6 |
|  | 264-412 | .0904 | 20.4 |
|  | 464-536 | .0143* | -3.3 |
| 6 | 264-476 | .0030* | -23.9 |
| 7 | 144-268 | .1168 | 26.6 |
|  | 328-484 | .2237 | -44.5 |
| 9 | -108-100 | .0023* | 34 |
|  | 304-580 | .0101* | 33.4 |
| 10 | 368-568 | .0000* | 36.5 |
| 11 | 336-488 | .0101* | 28.4 |
| 12 | 176-288 | .0395* | 39.5 |
|  | 320-556 | .0551 | 47.1 |
| 13 | 32-148 | .1165 | 34.3 |
|  | 396-496 | .1592 | -29.2 |

\*= statistically significant after FDR correction

**Figure 2:**
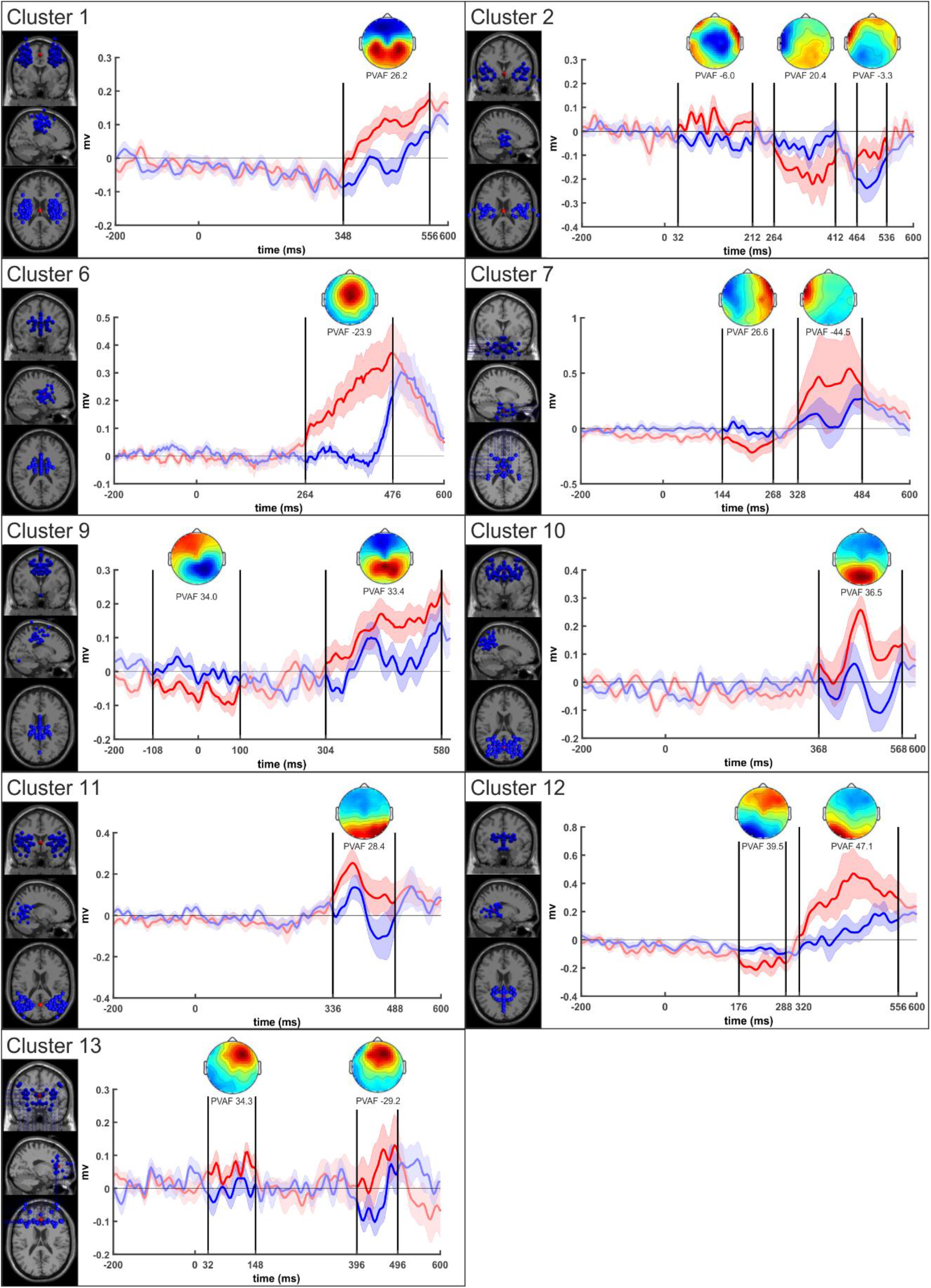
Clusters results. For each cluster are shown: the components’ dipoles’ location (on the left), the ERP waveform computed contrasting Aware (in red) and Unaware (in blue) conditions, the scalp distribution of the relative time-window and the percent variance accounted for (pvaf) each time-window. Time windows are highlighted by vertical black lines.

The same analyses were performed also on Correct-Incorrect comparison (i.e., Correct trials are those trials in which participants reported accurately the side of presentation of the stimulus). Correct trials represented on average the 79.13% of trials. As shown in Fig 1 (see Supplementary Material), the Correct vs Incorrect contrast yielded a grand average ERP very similar to the grand average ERP obtained contrasting Aware and Unaware epochs. The IC clustering, and the respective statistical analysis largely replicated the result of the Aware vs Unaware comparison (see, Supplementary Fig. 2).

## 4. Discussion

ERP correlates of conscious vision have identified two major correlates of visual awareness (VAN and LP). While scalp recorded ERPs display excellent time resolution, the sources of VAN and LP remain open. This is an important open question as theories of consciousness make different predictions about the location and the timing of consciousness-related activity in the brain (Seth & Bayne, 2022).

The present study aimed to characterize the neural dynamics underlying conscious visual perception by decomposing ERPs into independent components. This allowed us to identify and localize the sources of neural activity that contribute to the grand-average ERP correlates of conscious perception. Overall, in keeping with previous electrophysiological literature, the scalp recorded grand average ERPs obtained contrasting Aware and Unaware trials highlighted a significant difference in the N2 time-window (i.e., VAN), followed by a significant difference in the P3 amplitude (i.e., LP). Independent component analysis and clustering showed that activity of sources in many different cortical areas correlated with consciousness. In contrast to a serial, bottom-up driven process, the results suggest that the earliest differences between Aware and Unaware conditions were observed in parietal/frontal (cluster 9, between -108 and 100 ms) and temporal (cluster 2, between 32 and 212 ms) regions—that is, before stimulus-evoked activation. Also, early activity in prefrontal cortex correlated with conscious perception (cluster 13, between 32 and 148 ms), but this effect did not reach statistical significance. This early wave of conscious perception related activity was followed by correlates of conscious vision in occipito-temporal regions in a time-window corresponding to typical VAN time-window (cluster 2 until 212 ms, cluster 7, between 144 and 268 ms, cluster 12, between 176 and 288 ms). Finally, clusters spread over frontal, parietal, temporal and occipital areas displayed late differences in the P3 time-window. Notably, some of the sources that were active during LP time-window were active also during and before VAN (e.g., Cluster 2). Other sources were active solely during the LP (e.g., Cluster 10). This suggests that consciousness-related activity develops at least in part in “accumulative fashion” in a network of areas: consciousness-related activity in few early sources continues while additional sources are engaged as a function of time.

In general, our results are in accordance with previous source localization studies that identify the cortical generator of VAN in occipito-temporal brain regions (Vanni *et al*., 1997; Liu *et al*., 2012). Clusters 2, 7 and 12 revealed that the dipoles of components showing differences in the N2 amplitude when Aware and Unaware conditions were contrasted were localized in occipito-temporal areas. According to a popular interpretation, this activity in VAN time-window, possibly reflecting integrated, recurrent activity of multiple sources in the visual system, is the correlate of conscious visual perception (Förster *et al*., 2020; Mazzi *et al*., 2019; Dembski *et al*., 2021).

Dipoles of components reflecting differences in the LP amplitude were spread over frontal, parietal and occipito-temporal cortex, supporting the idea that LP has neural generators in wide-ranging cortical areas. According to a large body of literature, LP does not reflect neural processes purely related to subjective awareness of visual stimuli (Mazzi *et al*., 2020), but it is rather involved in later stages of processing such as processing task-relevant stimuli (Pitts *et al*., 2014; Shafto & Pitts, 2015), decision making (Koivisto & Grassini, 2016; Tagliabue *et al*., 2019), or processes related to reporting the contents of conscious perception (Koivisto *et al*., 2016). Our results are consistent with this interpretation as some LP sources were added “on top of” earlier consciousness-related sources. While this interpretation holds that conscious *perception* emerges in earlier time-windows (e.g., in VAN time-window), activity in LP could be related to higher-forms of conscious processing (e.g., whereas VAN may be sufficient for conscious perception of simple stimulus features, later activity may be related to consciously accessing semantic information about the perceived object; Jimenez et al., 2021).

Early activity observed in fronto-parietal areas in clusters 9 and 13 are so early (<150 ms after stimulus onset), that they are outside the time-windows typically considered to directly enable conscious perception. These components may therefore reflect top-down mechanisms that are “prerequisite” processes that occur before the stimulus enters the consciousness. Activity in fronto-parietal areas has been associated to visuospatial attention mechanisms (Corbetta et al., 2008; Parisi et al., 2020; Vossel et al., 2014). In particular, it has been proposed that attentional orienting towards specific locations is enabled by a bilateral fronto-parietal network, including the intraparietal sulcus (IPS), the superior parietal lobule (SPL) and the frontal eye fields (FEF) (Corbetta et al., 2000, 2008). Since in the present study participants were asked to report on which side of the screen the stimulus was presented, it is likely that they were covertly allocating their attention towards a location where the stimulus could appear. This allocation of attention may have helped to facilitate the entry of the visual input in visual awareness, without directly enabling conscious vision. That said, one could also argue that some of these early effects reflect proper conscious vision. While VAN latency is typically around 200 ms after stimulus onset, studies also show that VAN sometimes onsets around 100 ms (Koivisto *et al*., 2005, 2009; Koivisto & Revonsuo, 2008). Although visual attention modulates responses in the same time-window, the awareness related effect seems to emerge independently of attention (Koivisto et al., 2005; Koivisto et al., 2006). Also, in the present study the onset of scalp-recorded VAN was before 100 ms. Therefore, it remains possible that the earliest fronto-parietal clusters also contributed to early conscious perception. This possibility is intriguing, because it could indicate that fronto-parietal areas provide key top-down modulation which enable consciously accessing simple visual features rapidly (Railo et al., 2015). Arguably, the large diameter visual stimuli, and simple location detection task employed in the present study were key to observing the early VAN as these visual features may be efficiently processed, enabling rapid conscious perception (Kouider & Dehaene, 2007; Jimenez *et al*., 2021).

Although the IC-clustering method offers a promising approach to localize neural sources of the EEG signal, the method also has its limitations. First, because of the limited number of electrodes, lack of information about individual participants’ brain anatomy, and lack of information about the precise locations of electrodes in individual participants, the spatial resolution of the present source-localization is coarse. Second, ICA analysis and clustering is a statistical approach, and the results could be influenced by factors such as number of electrodes and clusters. Third, compared to the classical grand-average ERP correlates of conscious vision, the IC-correlates of conscious vision generally had smaller effect size. For these reasons, even though the sample size of the present study is larger than in most EEG studies of conscious vision, it could still be relatively limited for the proposed approach.

## 5. Conclusion

In the search for the NCCs, the present results provide further significant information about the spatio-temporal neural dynamics involved in conscious vision, highlighting that IC-clustering represents a useful tool to investigate the neural correlates of conscious perception. The novel approach adopted in the present study enabled us to unveil previously “hidden” sources of ERP correlates of conscious vision in widespread cortical areas. In addition to previously often reported correlates (VAN and LP), the results revealed earlier effects in fronto-parietal regions whose role in the emergence of visual awareness remains to be clarified.

## Acknowledgements

H.R. was funded by the Academy of Finland (grant <u>#308533</u>).

## Abbreviations

EEG: electroencephalography
ERP: event-related potential
ICA: independent component analysis
LP: Late Positivity
VAN: visual awareness negativity

## Data Accessibility Statement

The raw data supporting the conclusions of this article will be made available by the authors, without undue reservation, to any qualified researcher.

---

**SUPPLEMENTARY Figure 1.**
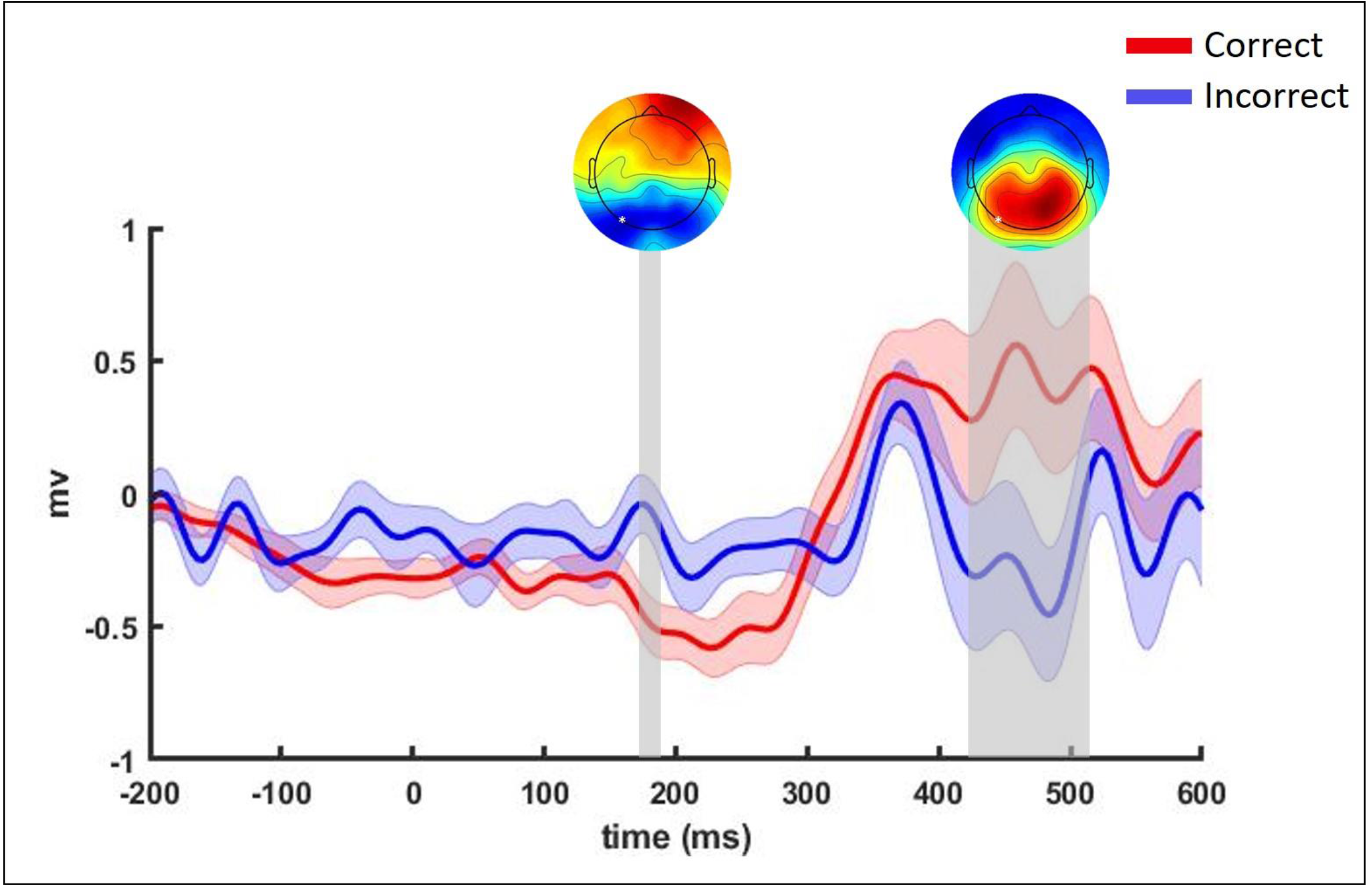
ERP results obtained contrasting Correct and Incorrect conditions. Grand-average of ERPs computed for Correct and Incorrect conditions at channel O1 (marked with a white star). Significant time-windows are highlighted in grey. The shaded area of the waveforms represents SEM at each time point, and scalp distribution maps represent the voltage difference between conditions

**SUPPLEMENTARY Figure 2:**
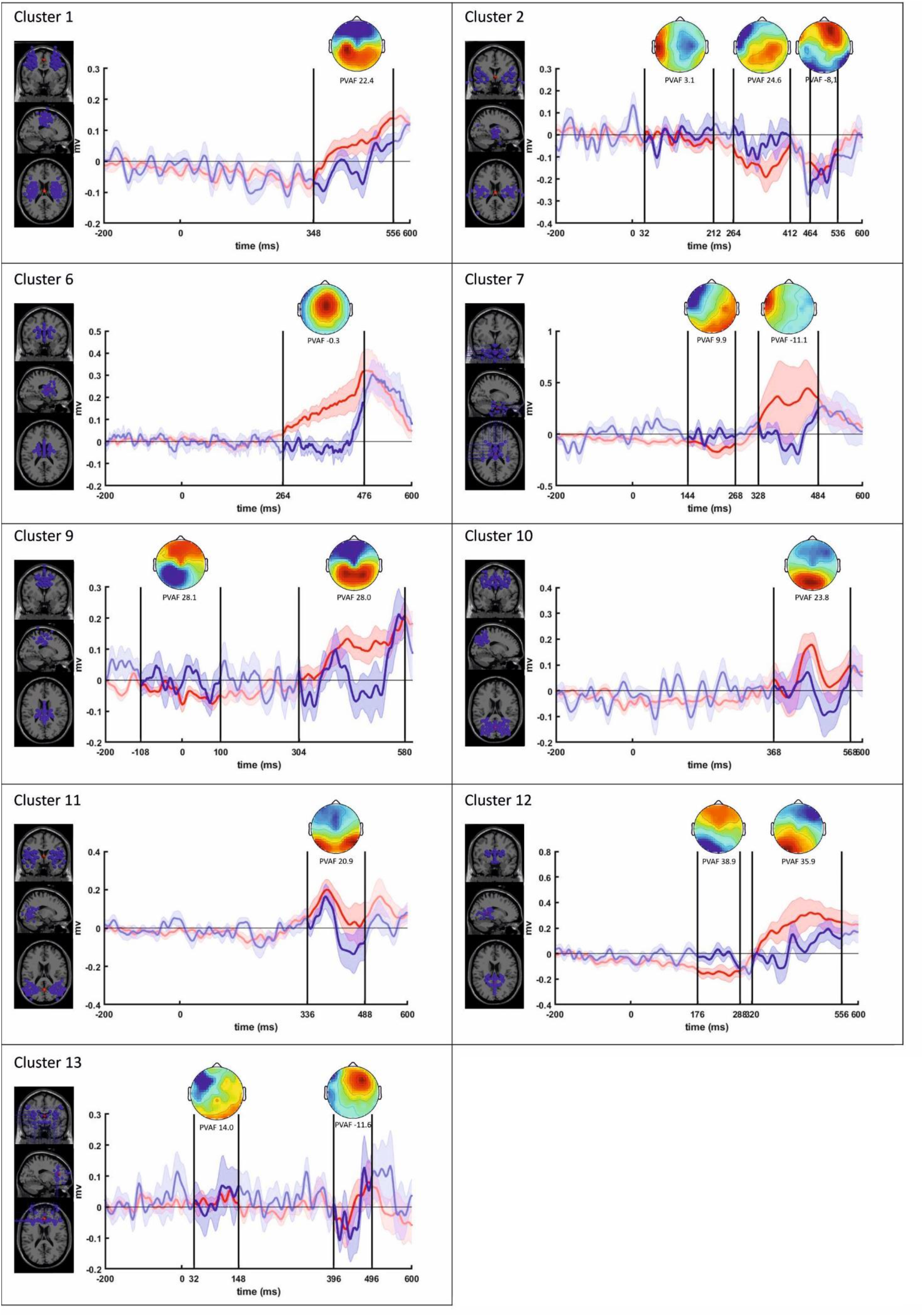
Clusters results. For each cluster are shown: the components’ dipoles’ location (on the left), the ERP waveform computed contrasting Correct (in red) and Incorrect (in blue) conditions, the scalp distribution of the relative time-window and the percent variance accounted for (pvaf) each time-window. Time windows are highlighted by vertical black lines.

**SUPPLEMENTARY Table 1:** Results of FDR-corrected paired-sample t-tests computed contrasting Correct and Incorrect conditions within each cluster in the reported time-windows.

| Cluster number | Temporal windows | p-value (fdr corrected) | Pvaf (%) |
| --- | --- | --- | --- |
| 1 | 348-556 | .023* | 22.4 |
| 2 | 32-212 | .762 | 3.1 |
|  | 264-412 | .165 | 24.6 |
|  | 464-536 | .317 | -8.1 |
| 6 | 264-476 | .023* | -0.3 |
| 7 | 144-268 | .285 | 9.9 |
|  | 328-484 | .283 | -11.1 |
| 9 | -108-100 | .258 | 28.1 |
|  | 304-580 | .023* | 28.0 |
| 10 | 368-568 | .031* | 23.8 |
| 11 | 336-488 | .023* | 20.9 |
| 12 | 176-288 | .023* | 38.9 |
|  | 320-556 | .057 | 35.9 |
| 13 | 32-148 | .967 | 14.0 |
|  | 396-496 | .879 | -11.6 |

